# Cisplatin is more mutagenic than carboplatin or oxaliplatin at equitoxic concentrations

**DOI:** 10.1101/2020.08.11.245969

**Authors:** Bernadett Szikriszt, Ádám Póti, Eszter Németh, Nnennaya Kanu, Charles Swanton, Dávid Szüts

## Abstract

Platinum-based drugs are a mainstay of cancer chemotherapy. However, their mutagenic effect can increase tumour heterogeneity, contribute to the evolution of treatment resistance, and also induce secondary malignancies. We coupled whole genome sequencing with phenotypic investigations on two cell line models to compare the magnitude and understand the mechanism of mutagenicity of cisplatin, carboplatin and oxaliplatin. Cisplatin induced significantly more base substitution mutations than carboplatin or oxaliplatin when used at equitoxic concentrations on human TK6 or chicken DT40 cells, and also induced the highest number of short insertions and deletions. Assessment through histone H2AX phosphorylation and single cell agarose gel electrophoresis suggested that cisplatin caused more DNA damage than carboplatin or oxaliplatin. The analysis of base substitution spectra revealed that all three tested platinum drugs elicit both a direct mutagenic effect at purine dinucleotides, and an indirect effect of accelerating endogenous mutagenic processes. Whereas the direct mutagenic effect correlated with the level of DNA damage caused, the indirect mutagenic effects were equal. The different mutagenicity and DNA damaging effect of equitoxic platinum drug treatments suggests that DNA damage independent mechanisms significantly contribute to their cytotoxicity. Thus, the comparatively high mutagenicity of cisplatin should be taken into account in the design of chemotherapeutic regimens.

## INTRODUCTION

The formation of new somatic mutations increases tumour heterogeneity and can contribute to treatment resistance. Therefore, there is much interest in assessing the mutagenicity of DNA-damaging cytotoxic treatments. Mutagenic chemotherapy can also contribute to the formation of secondary cancers through inducing mutations in normal somatic tissue.

The platinum drugs cisplatin, carboplatin and oxaliplatin are amongst the most commonly used cytotoxic agents, employed in the treatment of a wide range of cancer types. Cisplatin, which received US Food and Drug Administration (FDA) approval in 1978, is effective in treating e.g. bladder, head and neck, lung, ovarian, testicular cancer (1-5). Cisplatin is also a widely used agent in the treatment of paediatric tumours such as neuroblastoma, germ cell tumours, osteosarcoma, retinoblastoma, hepatoblastoma, brain tumours, spinal cord tumours and lymphomas (6). Cisplatin therapy is limited by serious side effects such as neurotoxicity, ototoxicity and nephrotoxicity, and the development of resistance (7-9). Carboplatin, an analogue of cisplatin selected for reduced nephrotoxicity, obtained FDA approval in 1989. Carboplatin is mainly used for ovarian, lung, and head and neck cancers (10-12). Neoadjuvant therapy based on either cisplatin or carboplatin has shown benefits in the treatment of triple negative breast cancer (13,14). Carboplatin is particularly effective compared to docetaxel in the management of germline mutant BRCA1/2 breast cancer (15).

Carboplatin is as effective as cisplatin in the treatment of ovarian cancer, but it is better tolerated (16), whereas it shows no clear tolerability benefit over cisplatin in the treatment of lung cancer (17). The third-generation platinum-derived antitumor drug oxaliplatin, which obtained FDA approval in 1996, is mainly used in colorectal cancer in combination therapy (18-20). Oxaliplatin has lower toxicity than cisplatin (21); it shows very limited activity against cisplatin-resistant tumours (22).

Platinum drugs damage DNA by forming covalent adducts. Cisplatin comprises a platinum atom surrounded in a plane by two ammonia groups and two chloride ligands. After entering the cell, the chloride ligands are sequentially displaced by water molecules, and the resulting positively charged platinum complexes can react with nucleophilic sites of the DNA (23,24). Cisplatin mainly reacts with purine bases, forming intrastrand crosslinks at GpG and ApG or GpGpG and ApGpG sequences (90% of all adducts), and rarer interstrand crosslinks (3-5%) (25,26). Cisplatin can also form DNA-platinum-protein complexes, for instance it can cross-link chromosomal proteins to DNA (27,28). Carboplatin possesses a similar chemical structure to cisplatin: a platinum atom surrounded in a plane by two ammonia groups and two leaving carboxylate groups in the *cis* position. The dicarboxylate unit of carboplatin hydrolyses much more slowly than the chloride groups of cisplatin (24), but the adduct-forming moiety is identical. Oxaliplatin has a similar dicarboxylate leaving group to carboplatin, but it also possesses a 1,2 diamino-cyclohexane ligand. The active form of oxaliplatin therefore differs from that of the previous two drugs, but it also forms DNA adducts in sequence- and region-specific manner. Oxaliplatin is less reactive with DNA compared to cisplatin (29), but nevertheless more cytotoxic (30).

The mutagenicity of cisplatin has been investigated and reported using prokaryotic and eukaryotic reporter gene assays (reviewed by 31,32) and by whole genome sequencing of animals or cell lines (33-36). The mutagenic footprint of platinum treatment was subsequently also found in the genomes of primary tumours (35,37,38) and metastases (39,40). In comparative studies, cisplatin and carboplatin were found to be similarly mutagenic in the Salmonella histidine reversion assay when a higher concentration of carboplatin was used (41), and cisplatin was found to be more mutagenic than oxaliplatin in the HPRT gene of mammalian cells at equimolar concentrations (42). Finally, 3.125 μM cisplatin was similarly mutagenic to 5 μM carboplatin on human induced pluripotent stem cells (36). In cancer genomes, somatic mutations attributed to each of the three discussed platinum agents showed varied contribution to the total mutational burden in different tissues (38).

To achieve a clinically relevant comparison of the mutagenicity of these three common platinum-containing cytotoxic agents, and to also analyse their mutation spectra, we assayed mutagenesis in the genomes of two different cultured cell lines upon treatments at carefully determined equitoxic concentrations. We were able to show that all three platinum drugs were mutagenic and induced both single nucleotide variations (SNVs) and short insertions/deletions (indels). Carboplatin and oxaliplatin caused fewer mutations than cisplatin, although induced similar mutational spectra that resembled the spectrum of cisplatin-induced mutations in cancer. Cisplatin caused more DNA damage at equitoxic concentration than carboplatin and oxaliplatin. The cause of the different mutagenic and DNA damaging effects of the tested platinum drugs is likely related to differences in their mechanisms of adduct formation and cell killing.

## RESULTS

### Platinum treatments at equitoxic concentrations

We employed two cell lines for testing the mutagenicity of platinum chemotherapeutics: the chicken DT40 lymphoblastoma cell line, which was used previously to establish the mutagenicity of cisplatin (34), and the human TK6 lymphoblastoid cell line, which has been extensively used for genotoxicity and mutagenicity assays (43). The karyotypic stability and low spontaneous mutation rate of DT40 makes this cell line especially suited for whole genome mutation detection (44). As the primary aim of this study was to directly compare the mutagenicity of platinum-containing therapeutic drugs, it was important to establish comparable treatment conditions. We opted for one-hour treatments, which is in the range of the plasma elimination times of free platinum drugs (45,46). We carefully measured the cytotoxicity of one-hour treatments with cisplatin, carboplatin and oxaliplatin using colony formation assays on DT40 cells, and cell viability measurements on TK6 cells (Figure 1). IC_75_ and IC_50_ concentrations, representing equitoxic doses that allow 75% or 50% cell survival, were determined using these curves and employed in all subsequent experiments (Table 1). The chosen treatment conditions showed good agreement between the relative toxicities of the three platinum drugs in DT40 and TK6 cells, with the TK6 cells showing slightly greater sensitivity. Also, the established treatment concentrations correlated very well with published pharmacokinetic measurements; the IC_50_ concentrations on the human TK6 cells were approximately 3-4 fold higher than the peak plasma concentrations of the respective platinum drugs (47-49).

**Table 1.**
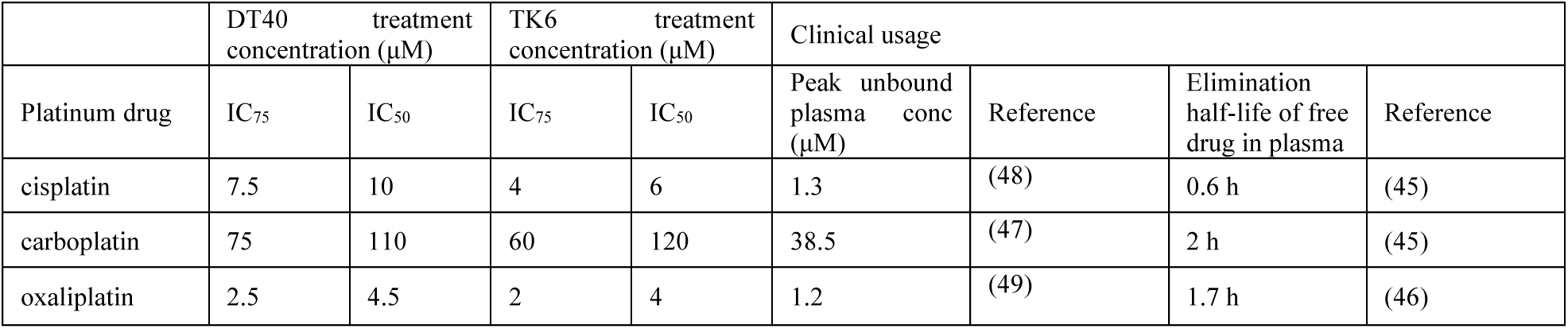
Treatment concentrations. Typical values from the literature are shown for comparison.

**Figure 1:**
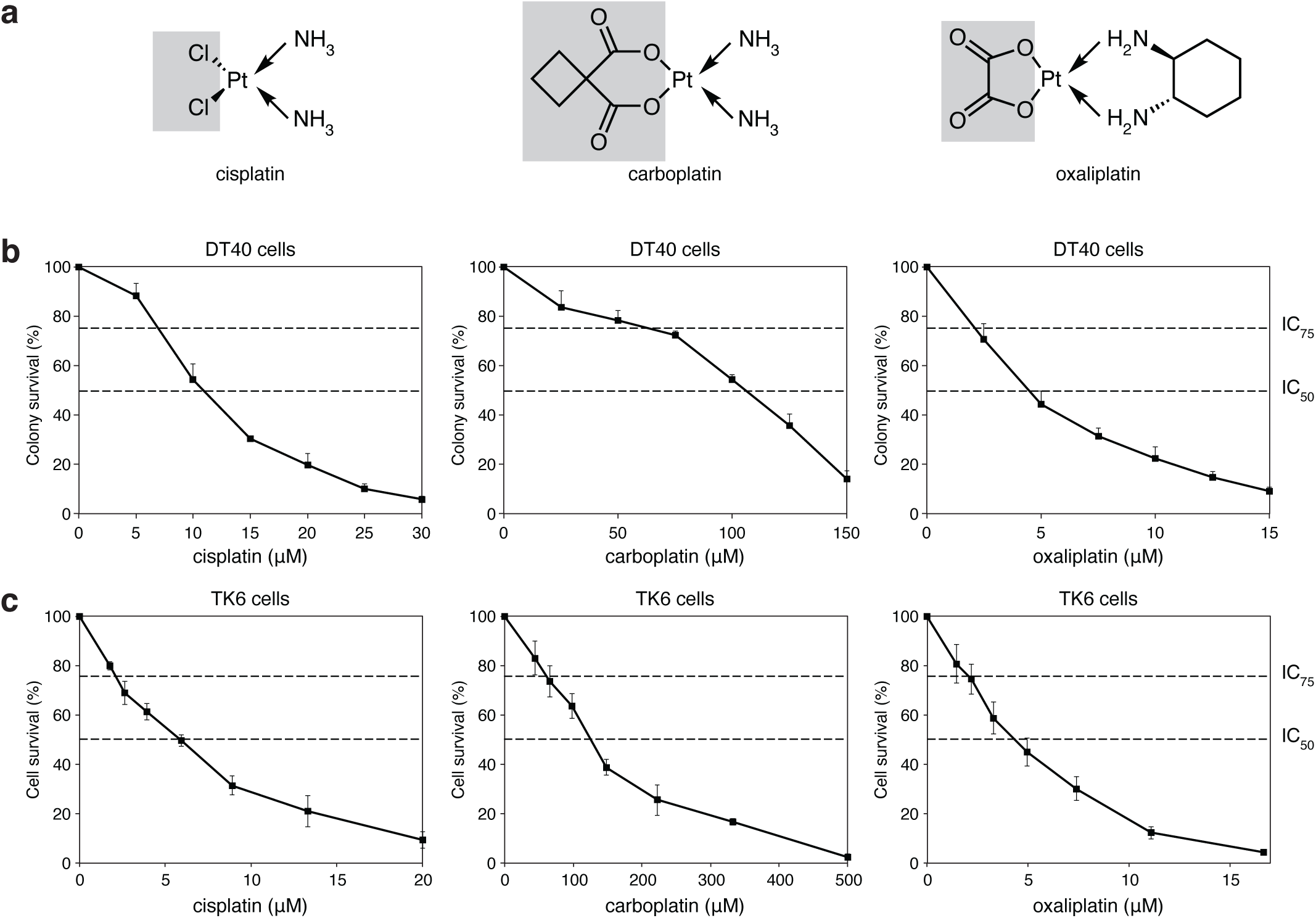
Platinum treatments. **a** Chemical structures of the investigated platinum drugs. The shaded parts of the molecules are removed prior to or during adduct formation. **b** Results of colony survival assays of DT40 cells treated with the indicated platinum drugs for one hour. **c** Results of cytotoxicity assay of TK6 cells treated with the indicated platinum drugs for one hour. The mean and S.E.M. of three independent experiments is shown in (**b, c)**; the concentrations chosen for the mutagenesis assays (IC_75_ and IC_50_ values) are indicated with dashed lines.

### Cisplatin is more mutagenic than carboplatin or oxaliplatin

To accurately determine and compare the mutagenicity of platinum agents, we submitted cell populations derived from single cell clones to one-hour treatments with each agent at IC_50_ or IC_75_ concentrations. We performed four repeated treatments at one-week intervals to mimic clinical conditions. We showed earlier that this treatment regimen at the IC_50_ concentration of cisplatin does not select for resistance (34). In parallel, we kept ‘mock-treated’ cells for the same overall duration. Whole genome sequences of the ancestral clones and 3-5 post-treatment descendant clones were compared with the IsoMut bioinformatic tool (50); any unique mutations that were detected in descendant clones must have arisen during the platinum or mock treatment.

Mutations were classified as single nucleotide variations (SNVs), short insertions and short deletions. This is the first study to determine the spontaneous mutation rates of the TK6 cell line, which were moderately higher than in DT40 cells, measured earlier as 0.46 per Gb diploid genome per cell cycle (34). Specifically, the mean spontaneous base substitution rate was 169 per diploid genome in 50 days, which is equivalent to 0.75 per Gb per cell cycle, calculating with a 16 h cell cycle time. This relatively low spontaneous mutation rate makes the TK6 cell line well suited for mutagenesis studies.

Compared to the mock treatment, the number of mutations significantly increased in all three categories in response to treatment with either platinum agent, in both DT40 and TK6 cells (Figure 2A, B, Supplementary Table S6, S11). The mutagenic effect of the platinum drugs was dose-dependent (Figure 2A). Importantly, cisplatin treatment was generally more mutagenic than carboplatin or oxaliplatin treatments at equitoxic concentrations. Specifically, cisplatin induced significantly more SNVs than carboplatin or oxaliplatin at either IC_75_ or IC_50_ concentrations in DT40 cells, and also induced significantly more insertions and deletions at most concentrations (Figure 2A). The SNV and insertion mutagenicity of carboplatin and oxaliplatin was similar at approximately half of that of cisplatin, or less. Interestingly, oxaliplatin induced significantly more short deletions than carboplatin (*p* = 0.008 at IC_50_, Student’s t-test), though other aspects of their mutagenicity were similar. The differences between the mutagenicity of platinum drugs were less significant in TK6 cells, but the trends for lower mutagenicity of carboplatin and oxaliplatin were also apparent (Figure 2B). In comparison to the mock treatment, cisplatin induced an average of 532 extra SNVs per TK6 genome, whereas carboplatin and oxaliplatin induced an average of 324 and 207 SNVs (61% and 39% of the cisplatin-induced SNVs, respectively).

**Figure 2:**
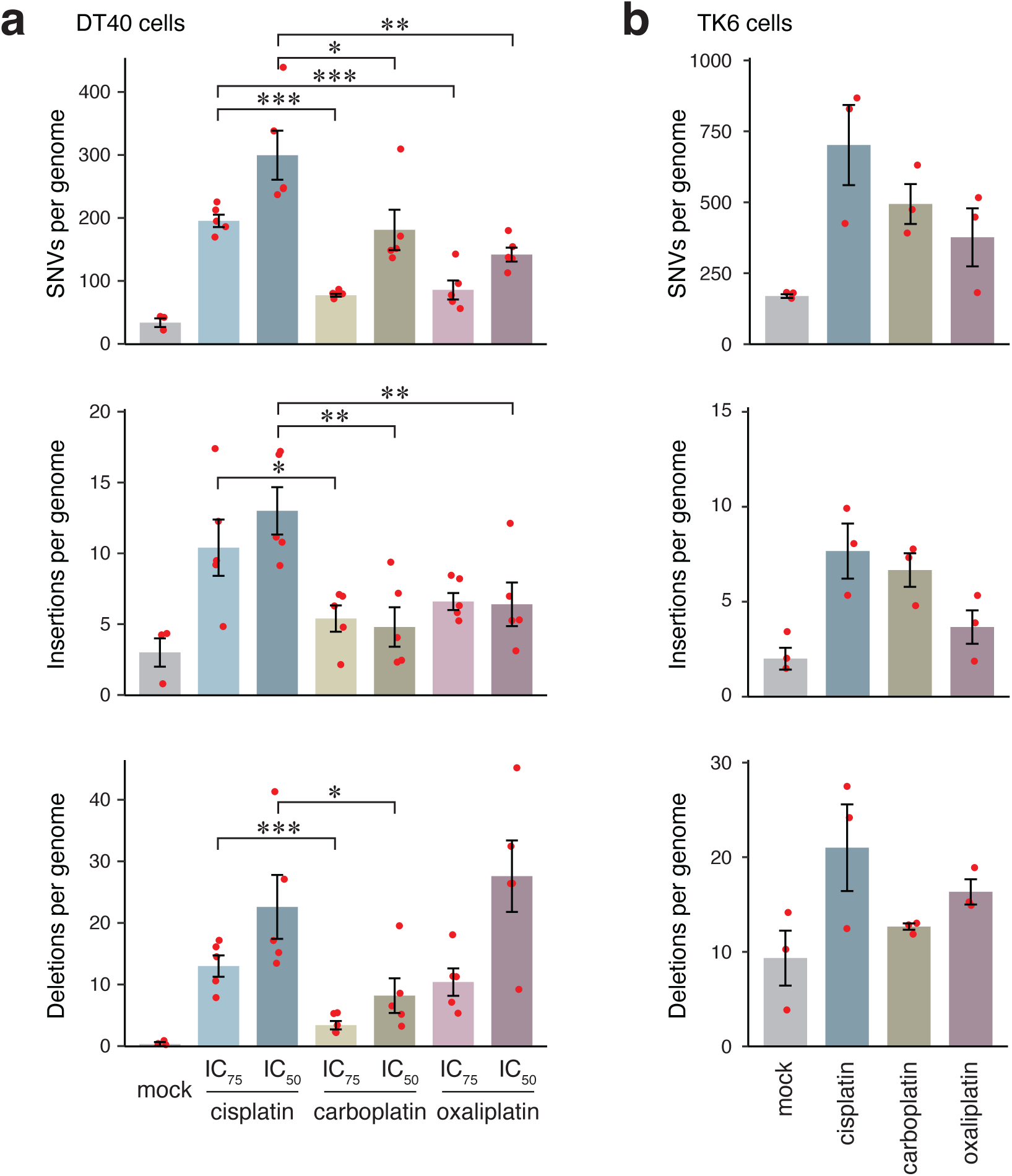
Platinum-induced mutagenesis. **a, b** The mean number of platinum-induced SNVs, short insertions and short deletions detected per sequenced genome in DT40 (**a**) and TK6 cell line (**b**) after mock treatment and the indicated platinum treatments. Red symbols show the values for individual samples, error bars indicate standard error of the mean (S.E.M.). Significance values are indicated (unpaired *t*-test, \**P*<0.05, \*\**P*<0.01, \*\*\**P*<0.001).

### Cisplatin induces more DNA damage than carboplatin or oxaliplatin

The different mutagenicity of the platinum drugs could be due to differences in DNA damaging activity, or differences in the mutagenicity of the repair of drug-induced lesions. To test the DNA damage caused by the treatments, we first assessed the accumulation of phosphorylated histone H2AX (γH2AX). H2AX is phosphorylated by ATM, ATR or DNA-PK, and serves as a marker for both DNA double strand breaks (DSBs) and for accumulated single stranded DNA at stalled replication forks (51,52). 24 hours after one-hour treatments at IC_75_ or IC_50_ concentrations, which were equivalent to the treatments used for the mutagenesis experiments, western blots showed an elevated level of γH2AX in whole cell extracts of DT40 cells in response to each platinum drug (Figure 3A). Cisplatin treatment induced significantly higher levels of γH2AX than either carboplatin or oxaliplatin at equitoxic levels. Treatment of TK6 cells with cisplatin also induced more γH2AX than carboplatin or oxaliplatin (Figure 3B). In a time-course following one-hour IC_50_ treatments, the drug-induced elevation of γH2AX levels was already observable two hours following the treatment in the case of all three drugs on both DT40 and TK6 cells (Figure 3C, D), suggesting that this is a direct effect due to the stalling and possible collapse of replication forks rather than to DNA breaks caused by apoptosis. γH2AX accumulation in subnuclear foci 24 h after platinum treatment was also apparent using immunofluorescence (Figure 3E, G). The number of γH2AX foci was in general higher upon cisplatin treatment than upon treatment with carboplatin or oxaliplatin in both DT40 and TK6 cells, at either IC_75_ or IC_50_ concentrations (Figure 3F, H). The γH2AX focus counts confirmed the western blot results that cisplatin induces more damage than the other tested drugs at equitoxic concentration.

**Figure 3:**
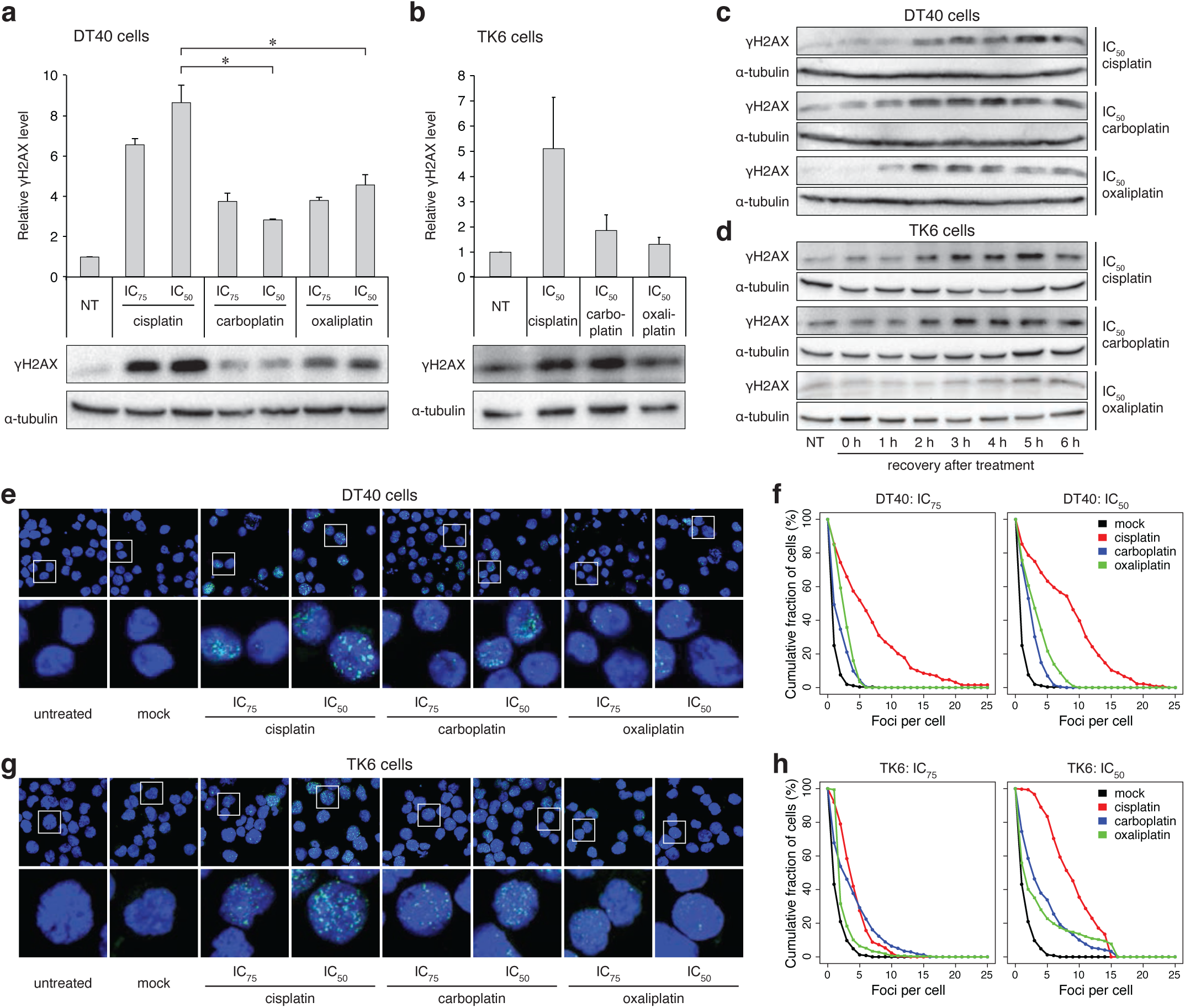
Markers of DNA damage response. **a, b** Western blots detecting γH2AX in DT40 cells (**a**) and TK6 cells (**b**); subjected to no treatment (NT) or to 1 h treatment with IC_75_ and IC_50_ cisplatin, carboplatin and oxaliplatin with a 24 h recovery period. Above the representative chemiluminescence images, mean values of normalised signals from *n*=3 experiments are shown. Error bars indicate S.E.M. Significant differences of values between cisplatin and other platinum treatments are indicated (paired *t*-test, \**P*<0.05). **c, d** Representative chemiluminescence images of γH2AX detected in DT40 cells (**c**) or TK6 cells (**d**) subjected to no treatment (NT) or to 1 h treatment at IC_50_ concentration with cisplatin, carboplatin or oxaliplatin, followed by 0, 1, 2, 3, 4, 5 and 6 h recovery period. Different exposure times were used to aid γH2AX visualisation. **e** Representative immunofluorescence images of γH2AX (green) and DNA (blue) in nuclei of DT40 cells after 1 h treatment with IC_75_ and IC_50_ cisplatin, carboplatin and oxaliplatin and a 24 h recovery period. **f** Proportion of DT40 cells in which at least the indicated number of γH2AX foci were detected after treatments as in (**e**). **g** Representative immunofluorescence images of γH2AX (green) and DNA (blue) in nuclei of TK6 cells after 1 h treatment with IC_75_ and IC_50_ cisplatin, carboplatin and oxaliplatin and a 24 h recovery period. **h** Proportion of TK6 cells in which at least the indicated number of γH2AX foci were detected after treatments as in (**g**).

We also measured DSB formation directly using single cell agarose electrophoresis comet assays. We observed a noticeable formation of comet tails following treatment with platinum agents at IC_50_ concentrations, although the signal was much weaker than that obtained with the positive control hydrogen peroxide, used at a sublethal concentration (Figure 4A, B). The quantification of tail moments revealed that cisplatin caused more excessive DNA breakage than carboplatin or oxaliplatin in both DT40 and TK6 cells (Figure 4C, D), which may be a direct effect from collapsing replication forks or the consequence of an earlier start of apoptosis in cisplatin-treated cells. In agreement with the induction of DNA breaks, cell cycle analysis using flow cytometry on DT40 cells showed an increased G2/M phase population, an increase in cells arrested in S phase, and an increase in apoptotic cells with a sub-G1 DNA content (Figure 4E, F). The G2 arrest and the apoptotic enrichment were more pronounced after cisplatin treatment, especially 16 h after treatment, though by 24 h the oxaliplatin-treated cells also started entering apoptosis. Taken together, these results highlight a correlation between the mutagenic and the DNA break-inducing effect of the various platinum agents.

**Figure 4:**
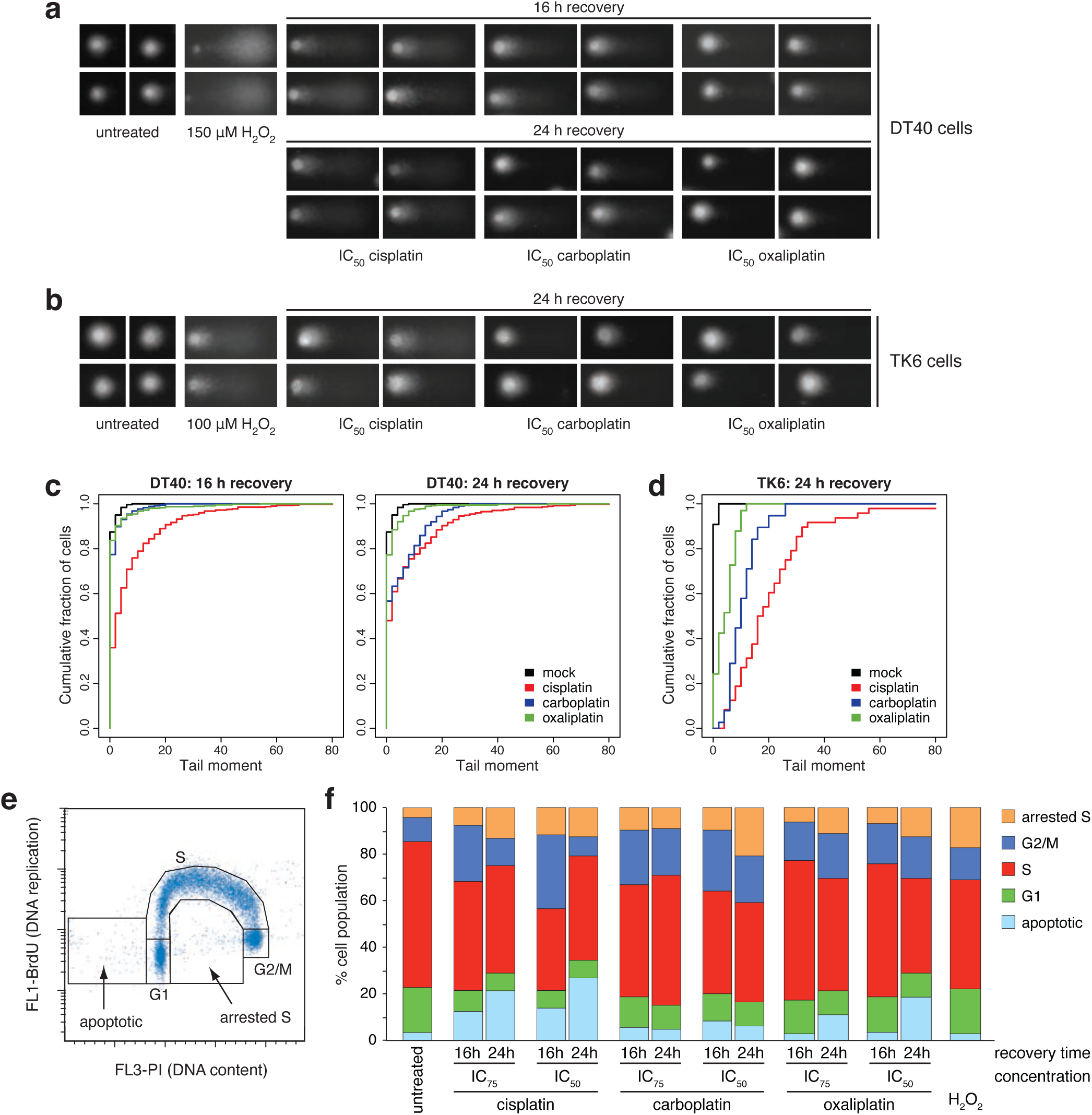
DNA breaks and cell cycle changes following platinum treatment. **a** Representative images of alkaline single cell gel electrophoresis (comet) assay 16 h and 24 h after 1 h IC_50_ cisplatin, carboplatin and oxaliplatin treatments in DT40 cells. **b** Representative comet images 24 h after 1 h IC_50_ cisplatin, carboplatin and oxaliplatin treatments in TK6 cells. 20 min treatments with 150 µM H_2_O_2_ serve as positive control in (**a, b**) **c, d** Proportion of the cells with at least the indicated tail moment values in DT40 cells (**c**) and TK6 cells (**d**). The tail moment values were calculated using the *CometScore 2*.*0* software. **e, f** Cell cycle analysis. The DNA content of DT40 cells was measured using propidium iodide staining (horizontal axis), and the rate of DNA replication using anti-BrdU-FITC antibody staining (vertical axis). **e** Rationale for assigning cell populations to different cell cycle phases, an apoptotic category with sub-G1 DNA content, and an arrested S phase category for non-replicating cells with an S phase DNA content. **f** The percentage of cells in each category. Cell cycle distributions were measured in untreated cells or after 1 h IC_50_ and IC_75_ treatments with the indicated platinum drugs followed by a 16 h or 24 h recovery period. 20 min treatments with 100 µM H_2_O_2_ served as a control. The mean of three independent experiments is shown.

### Platinum-induced mutation spectra reveal direct and indirect mutagenic effects

When base substitutions are viewed in the context of the neighbouring bases, the DT40-derived triplet mutation spectra of all three tested platinum agents are dominated by specific C>A and T>A peaks (Figure 5A). The common cisplatin-induced NCC>NAC mutations likely formed at intrastrand platinum crosslinks at GG dinucleotides, in agreement with the conclusions of our previous study (34). We also observed specific T>A substitutions and an increase in most triplet base substitution types. The mutation spectra of carboplatin- and oxaliplatin-treated DT40 cells showed similarities with the cisplatin spectrum, but the specific peaks were less prominent (Figure 5A). C>A and T>A mutations more commonly occurred on the transcribed strand, suggesting transcription-coupled repair of GG and AG intrastrand crosslink adducts (Supplementary Figure 1). Even though the platinum drugs induced a greater number of base substitutions in the TK6 cell genomes, the resulting spectra were relatively featureless (Supplementary Figure 2), although some similarities to the DT40-derived spectra such as the emergence of CTN>CAN mutations were apparent.

**Figure 5:**
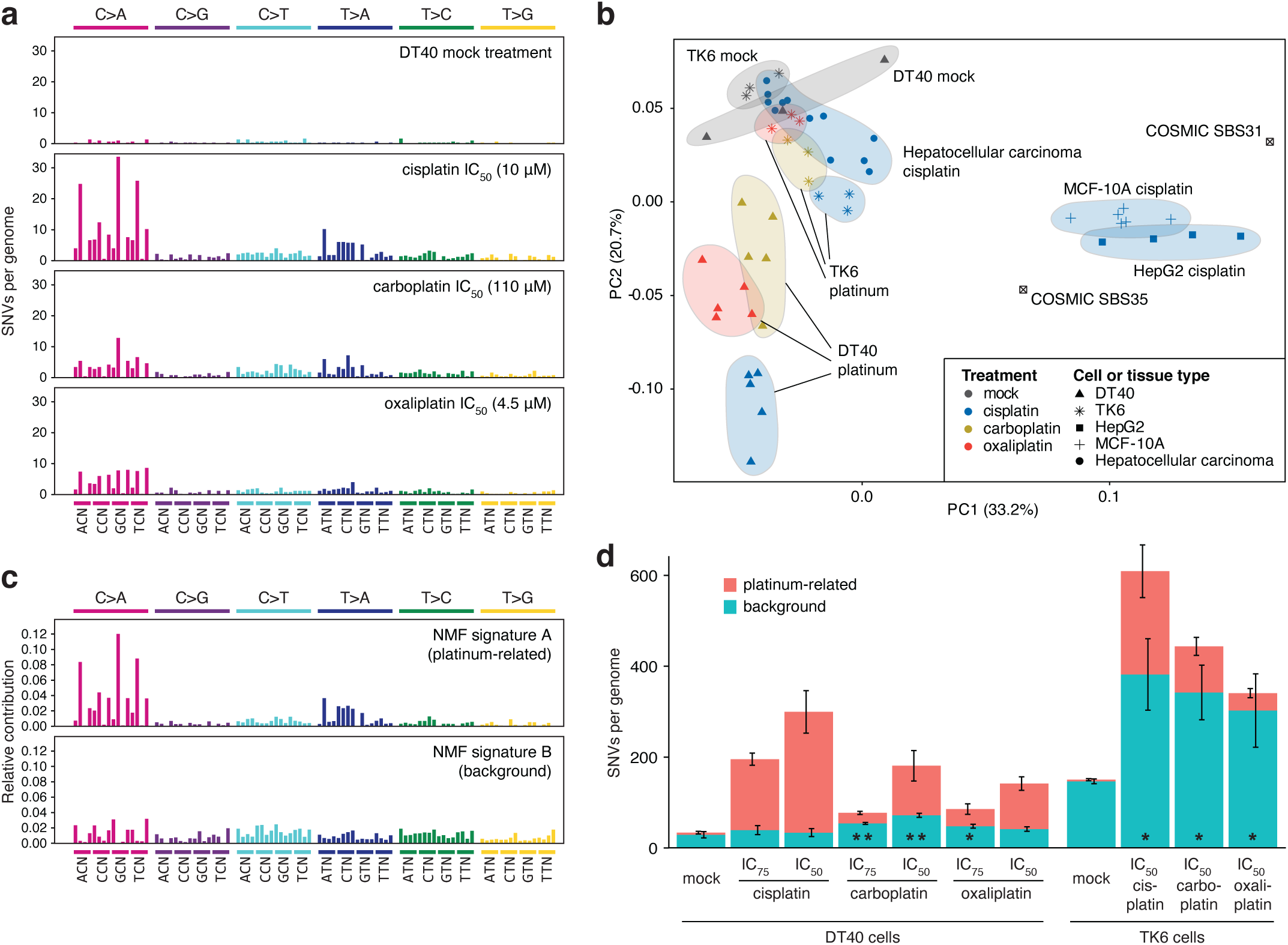
Analysis of SNV mutation spectra. **a** Triplet mutation spectra of the mock, IC_50_ (10 µM) cisplatin, (110 µM) carboplatin and (4,5 µM) oxaliplatin treatments of DT40 cells. The middle base of each triplet, listed at the *bottom*, mutated as indicated at the *top of the panel*. The third base of the triplets, not shown for lack of space, is alphabetical. **b** Principle component analysis of normalised triplet SNV spectra from all sampled sequenced in this study, as well as from cisplatin-treated MCF-10A and HepG2 cells (35) and of 12 cisplatin-treated hepatocellular carcinoma samples. The symbols indicate cell or tissue types, the colours specify platinum drug treatments. COSMIC signatures SBS31 and SBS35 were also included and are shown as labelled. **c, d** De novo NMF of SNV mutation spectra of all sequenced samples identified two components; triplet mutation spectra of NMF signature A (platinum related) and NMF signature B (background) are shown as the contribution of each triplet mutation type. **d** Mean SNV counts for all sequenced DT40 and in TK6 genomes, split using NMF into ‘platinum related’ and ‘background’ signatures and then averaged by cell type and treatment as indicated. The error bars indicate the (S.E.M) of the column below, based on five DT40 or three TK6 independent treated single cell clones. Significant differences in the mean number of mutations attributed to the ‘background’ signature, as compared to the mock treatment, are indicated at the bottom of the columns (unpaired *t*-test, \**P*<0.05, \*\**P*<0.01).

In order to better understand the base substitution processes upon platinum treatment, we performed principal component analysis (PCA) on the experimental triplet spectra. We included mutation datasets of cisplatin-treated human cell lines and of tumours in which the effect of cisplatin-induced mutagenesis was detected, taken from a recent publication (35). The results were intriguing (Figure 5B). In DT40 cells, principal component 2 (PC2) correlated very well with platinum-induced base substitutions. A similarly good correlation between the PCA result and mutagenicity was apparent in the case of TK6 genomes, except that here a combination of PC1 and PC2 contributes to the result. The analysis of both cell lines suggested that the tested platinum drugs cause base substitutions by similar mechanisms, and the only important difference is the level of mutagenesis. This conclusion is supported by the strong similarity of the dinucleotide mutation spectra and the types and context of short indels induced by the three drugs (Supplementary Figures 3-6). The mutation spectrum of cisplatin-treated MCF-10A and HepG2 cells, with many C>T mutations (35), seemingly better correlates with PC1 (Figure 5B). This suggests that there are cell type-dependent differences between platinum-induced mutagenic spectra, with the human TK6 cell line spectrum falling between chicken DT40 cells and human MCF-10A and HepG2 cells. The somatic mutation spectra of cisplatin-treated hepatocellular carcinomas most closely resembled that of TK6 cells (Figure 5B), therefore this cell line appears to be a good model for platinum mutagenesis in human cancer.

We attempted to extract the base substitution processes from our cell line dataset using non-negative matrix factorisation (NMF). NMF on all the DT40 and TK6 mutation data could fit two components with small error (Figure 5C, D; Supplementary Tables 12-14). One NMF component was only detected in platinum-treated samples, while the other component described spontaneous mutagenesis in mock treated DT40 and TK6 cells. The spectrum of the ‘background’ component resembled that of the ‘featureless’ COSMIC signatures SBS3, SBS5 and SBS40, of which SBS5 and SBS40 show correlation with the patient’s age in some cancer types, and are thought to represent mutagenesis in normal somatic cells (37,53). Unexpectedly, upon platinum treatment we observed a strong increase in the number of mutations that belong to this background signature, as compared to the matched mock treatment (Figure 5D). The increase was significant in TK6 cells upon treatment with any of the platinum agents. When we restricted the NMF analysis to the DT40 samples, the increase in ‘background’-type mutagenesis was also significant with all three drugs in these genomes (Supplementary Figure S2C). These novel findings suggest that in addition to causing mutagenic DNA adducts, platinum drugs also increase mutagenesis indirectly by accelerating the spontaneous mutagenic processes in treated cells, and the indirect mutagenic effect may even be the major contributor to the mutagenic effect of platinum treatment.

## DISCUSSION

In this study, we used two cell line-based mutagenesis models to investigate and compare the mutagenic effect of three commonly used platinum-based cytotoxic drugs. Whole genome sequencing revealed dose-dependent mutagenic effects of each tested drug. Importantly, cisplatin was significantly more mutagenic than carboplatin or oxaliplatin and also caused higher levels of DNA damage at concentrations of equivalent toxicity. The analysis of mutation spectra suggested that in addition to a direct mutagenic effect resulting from intrastrand platinum adducts, the drugs also damage DNA indirectly through a non-specific effect that may be correlated with their toxicity.

Whole genome sequencing provides the ultimate tool for determining the mutagenicity of environmental insults, including natural mutagens such as UV radiation (54), and mutagenic drugs (34). The correlation of cancer-derived mutational signatures with such environmental impacts can help with establishing their mutagenic effects, but the analysis is hampered by different exposures, different genetic backgrounds and other confounders. Only controlled experimental settings with single-parameter differences between treated and control samples can give reliable information on the full mutagenic effect of treatments. Whenever available, it is therefore best to use experimentally obtained mutation spectra for deciphering the biological mechanisms of mutagenesis. Our results demonstrated that platinum drugs induce mutations with two distinct spectra, suggesting a direct and an indirect mutagenic mechanism. The direct mechanism gives rise to point mutations and short indels mostly at GG and AG dinucleotides, presumably sites of platinum-containing intrastrand purine-purine crosslinks (26), and the resulting specific mutation types have also been observed in the genomes of platinum-treated tumours. When comparing the ‘direct’ platinum-related mutational signature derived from our data to platinum-associated COSMIC signatures, we can spot close similarities to SBS35 with identical prominent C>A and T>A mutation types, but limited likeness to SBS31 that is dominated by C>T mutations. SBS35 appears to describe platinum-induced base substitution mutations in cancer better than SBS31 (37), whereas SBS31 resembles the spectra derived from two other cisplatin-treated cell lines (35). Establishing the causes of the different balance of C>A, C>T and T>A mutations in various platinum-treated cell line models and cancers will require further investigation. We inferred a second, indirect mechanism from the increased contribution of a broad-spectrum SNV signature to overall mutagenesis. As this signature matches the mutational spectrum of mock treated control cells, its presence is not specific to platinum treatment. Therefore, the observed indirect platinum-induced mutational process would be difficult to detect in tumour genomes, supporting our cell line-based comparative approach. We can only speculate as to the cause of the platinum-induced ‘background signature’ mutations, which echo our previous observation of increased broad-spectrum mutagenesis upon etoposide treatment (34). The underlying mutagenic process also appears to operate in untreated cells and is therefore probably related to endogenous DNA damage, although we have little understanding of the mechanism of spontaneous mutagenic processes. Platinum treatment stressing the cells may enhance this effect by increasing the production of reactive oxygen species and other adduct-forming species, by reducing the fidelity of DNA synthesis by disturbing intracellular nucleotide availability, or by reducing the general efficiency of DNA repair.

The amount of treatment-induced DNA damage, indicated primarily by the γH2AX signal that appeared soon after treatment, correlated with the mutagenicity of the three tested drugs. Cisplatin appeared to cause more DNA damage and more mutations than carboplatin and oxaliplatin, and the extra mutations were primarily of the direct type at the site of lesions. It follows that at equitoxic concentrations cisplatin causes more DNA adducts. Successful replicative bypass of these lesions then gives rise to the observed mutations, while the collapse of some stalled replication forks at lesion sites can cause DNA breaks. But how could the matched treatments have equal toxicity, if cisplatin caused more DNA damage? It is important to note that platinum drugs form both intrastrand and interstrand DNA adducts, of which only the more common intrastrand adducts contribute significantly to mutagenesis (34), even though interstrand adducts may be more toxic. However, the ratios of various DNA adducts was found to be similar in the case of cisplatin and oxaliplatin (55). As platinum drugs readily react with proteins as well as nucleic acids, it is likely that their mechanism of cell killing also includes DNA-independent routes. Indeed, one such mechanism has been demonstrated for oxaliplatin, which exerts a cytotoxic effect via inducing ribosome biogenesis stress (56). The platinum agents may also stress cells via further, yet undiscovered mechanisms, as implied by the increased rate of background mutagenesis upon treatment with all three drugs. The differences may be due to the different relative access of the platinum drugs to DNA versus other cellular components. This could be related to their different rates of activation after cellular import (replacement of the chloride or carboxyl moieties with hydroxyl groups), and the different reactivity of the active forms. It is important to note that although cisplatin and carboplatin form the same *cis*-diammineplatinum crosslink adduct, their active forms and the initial adduct-forming reactions are different, as cisplatin only loses one of its chloride ligands prior to forming the first covalent bond with DNA (57), and carboplatin may react in an analogous sequential manner.

Differences in the mutagenicity of cytotoxic drugs are relevant for treatment-induced secondary cancers. The carcinogenicity of cisplatin-based therapy, recently reviewed by Liang and colleagues (58), is well documented: there is an increased risk of developing both solid tumours and leukaemia following cisplatin treatment of testicular or ovarian cancer. The appearance of secondary malignancies is a particularly significant issue in the treatment of childhood cancer (59). Our results suggest that the replacement of cisplatin with carboplatin could reduce the carcinogenicity of therapies. Mutagenic treatments may also accelerate the evolution of resistance in the primary tumours, as demonstrated by cisplatin-induced reversion mutations in *BRCA2* deficient cancers (34,60), especially if DNA repair deficiencies in the tumours increase the mutagenicity of the treatment (61). The genomic imprint of defective homologous recombination is a positive predictor of platinum treatment in advanced or triple negative breast cancer (14,62). In these cases, the choice of cisplatin or carboplatin may influence the evolution of resistance.

Further clinical relevance of our findings may concern the dose-limiting side effects of chemotherapeutic treatments. Platinum treatments show considerable peripheral neurotoxicity, which is much stronger in the case of cisplatin than carboplatin. The main proposed underlying mechanisms are nuclear DNA damage in dorsal root ganglia, or mitochondrial damage and oxidative stress (63). Our results suggest the dominance of the former mechanism based on greater nuclear DNA damage caused by cisplatin treatment. In contrast, the high nephrotoxicity of cisplatin could be independent of its DNA damaging effect and rather be related to the reactivity of this halogenated compound with cellular thiols (64). It will require further studies to establish whether the indirect mutagenic effect of all three platinum drugs observed in this study is related to oxidative stress, and whether this phenomenon is related to any specific side effects.

In conclusion, this genomic study demonstrated greater mutagenicity of cisplatin compared to carboplatin and oxaliplatin. We showed that platinum drugs exert direct as well as indirect mutagenic effects, of which the direct mutagenic effects correlate with the amount of DNA damage caused by the treatments but not with cytotoxicity. Our results can contribute to a careful appraisal of the benefits versus the short-term and long-term side effects of platinum-containing chemotherapeutics to guide therapeutic choices.

## METHODS

### Cell culture, platinum drugs

The wild type chicken DT40 cell line used in this study has been described (44). The human TK6 cell line was obtained from the TK6 mutants consortium (http://www.nihs.go.jp/dgm/tk6.html) (43). DT40 cells were grown in RPMI-1640 medium (Gibco, Life Technologies) supplemented with 7% fetal bovine serum (FBS) (Gibco, Life Technologies), 3% chicken serum (Sigma), 1% penicillin-streptomycin (Lonza) and 50 μM β-mercaptoethanol. TK6 cells were also maintained in the RPMI-1640 medium completed with 10% FBS and 1% penicillin/streptomycin. Both cell lines were cultured at 37 °C under 5% CO_2_. Cisplatin, carboplatin and oxaliplatin were obtained from Sigma-Aldrich.

### Platinum sensitivity assays and long-term treatments for mutagenesis experiments

Sensitivity to cisplatin, carboplatin and oxaliplatin was measured using colony survival assays on DT40 cells and cytotoxicity assays on TK6 cells. Ten thousand cells were treated for one hour at different drug concentrations. After the treatments, the platinum drug-containing medium was removed. For the colony formation assay the cells were plated onto 6-well plates in medium containing 1% methylcellulose using a ten-fold dilution series. After 10–12 days the surviving colonies were counted. For cytotoxicity assays, treated cells were plated onto 96-well plates at a density of 5000 cells/well, and cell survival was measured 72 hours later on a Perkin-Elmer EnSpire plate reader using the PrestoBlue cell viability reagent (Invitrogen).

For the mutagenesis experiment, single ancestral clones were isolated from the DT40 and TK6 wild type cell lines and expanded, then four rounds of drug treatments were performed in weekly intervals. In each treatment one million cells were exposed for one hour. Each treatment series lasted 50 days including the initial period of ancestral clone expansion. Mock treated cells were handled in parallel with the treated cells without the addition of platinum drug. Following the fourth treatment, single cell clones were isolated and expanded to one million cells prior to genomic DNA preparation using the Gentra Puregene Cell Kit (Qiagen).

### Whole genome sequencing, mutation calling and mutation analysis

Library preparation and DNA sequencing was done at Novogene, Beijing, using Illumina HiSeq X Ten and Novaseq instruments (2×150 bp PE) and at BGI, Hong Kong, using 2×100 bp BGISeq and 2×150 bp DNBSeq technology. A mean sequence coverage of 30x was targeted in all cases (Supplementary Tables S1, S7). DT40 sequence reads were aligned to the chicken (*Gallus gallus*) reference genome Galgal4.73; TK6 sequence reads were aligned to the human reference sequence. Duplicate reads were removed and the aligned reads were realigned near indels as described (61).

For detection of the independent SNVs and indels we used the IsoMut method developed for the detection of unique mutations in multiple isogenic samples (50). Briefly, after running IsoMut with default parameters, detected mutations were filtered using a probability-based quality score calculated from the mutated sample and one other sample with the lowest reference allele frequency. The quality score threshold was set such that no more than five false positive SNVs or one insertion or one deletion would be detected in pre-treatment starting clones. Detailed lists of detected mutations are presented in Supplementary Tables 2-6, 8-11. Individual SNV spectra were averaged for each treatment. *De novo* NMF decomposition and analysis of transcriptional strand bias was performed using the R package *MutationalPatterns* (65). Custom scripts are available at https://github.com/szutsgroup/platinum_mutagenesis.

### Detection of DNA damage markers

For western blot analyses, whole cell extracts were separated by sodium dodecyl sulphate-polyacrylamide gel electrophoresis and transferred to polyvinylidene difluoride membranes. After blocking in Tris-buffered saline (TBS) containing 3% bovine serum albumin for 1 hour, the membranes were incubated with anti-phospho-histone H2AX (Ser139), clone JBW301 primary antibody (Millipore, 05-636) in 1:2000 dilution at 4°C overnight, washed, and incubated with anti-mouse IgG-peroxidase secondary antibody (Sigma, A9044) in 1:20000 dilution for 1 h at room temperature. Chemiluminescent imaging using the Clarity Western ECL Blotting Substrate was performed with a ChemiDoc MP system and the band intensities were quantified using the ImageLab software (all from Bio-Rad Laboratories). Band intensities were normalised to α-tubulin detected on the same membrane with primary antibody T6199 (Sigma) at 1:500 dilution. Before averaging of measurements, the signals were also normalised to the untreated samples.

For immunofluorescence analysis, 1 million treated and recovered cells were pelleted onto poly-L-lysine (Sigma) coated coverslips and fixed with 4% paraformaldehyde in PBS. The samples were blocked with 0.1% Tween 20 and 0.02% SDS in phosphate-buffered saline (PBS), then incubated with anti-phospho-histone H2AX primary antibody in 1:1000 dilution and Alexa Fluor 488 anti-mouse secondary antibody (Thermo Fisher Scientific) in 1:1000 dilution for 1 hour each at 37°C followed by Hoechst 33342 (Thermo Fisher Scientific) in 1:10000 dilution at room temperature for 10 minutes. A Zeiss LSM710 confocal microscope was used to detect the fluorescent signal.

### Alkaline single cell gel electrophoresis

For alkaline single cell gel electrophoresis 0.5 million DT40 and TK6 cells were treated with platinum drugs at IC_50_ and IC_75_ concentrations for one hour and then allowed to recover for various periods. Samples were prepared using the Trevigen Comet Assay Kit, following the manufacturer’s protocols and recommendations. After the alkaline gel electrophoresis the samples were stained with 10 μg/ml propidium-iodide for 30 minutes at room temperature and viewed the with a LEICA DM IL LED fluorescence microscope. The CometScore 2.0 software was used for data analysis.

### 2D cell cycle analysis and flow cytometry

DT40 cells were treated with cisplatin, carboplatin or oxaliplatin for 1 hour and allowed to recover for 16 or 24 hours. Recovered treated cells were labelled with bromodeoxyuridine (BrdU), fixed and prepared for cell cycle analysis as described (66). The samples were analysed using flow cytometry on an Attune NxT flow cytometer (Thermo Fisher Scientific).

## DATA AVAILABILITY

Raw sequence data has been deposited with the European Nucleotide Archive under study accession number XXXX. [Submission in progress]

## DECLARATIONS

### Conflict of interest statement

C.S. receives grant support from Pfizer, AstraZeneca, BMS, Roche-Ventana, Boehringer-Ingelheim and Ono Pharmaceutical. C.S. has consulted for Pfizer, Novartis, GlaxoSmithKline, MSD, BMS, Celgene, AstraZeneca, Illumina, Genentech, Roche-Ventana, GRAIL, Medicxi, and the Sarah Cannon Research Institute. C.S. is a shareholder of Apogen Biotechnologies, Epic Bioscience, GRAIL, and has stock options in and is co-founder of Achilles Therapeutics. D.S. has consulted for Turbine Simulated Cell Technologies.

### Funding

This work was supported by the National Research, Development and Innovation Fund of Hungary (PD_121381 to BS and DS., K_124881 to DS, FIEK_16-1-2016-0005 to DS. CS is Royal Society Napier Research Professor. This work was supported by the Francis Crick Institute that receives its core funding from Cancer Research UK (FC001169, FC001202), the UK Medical Research Council (FC001169, FC001202), and the Wellcome Trust (FC001169, FC001202). CS is funded by Cancer Research UK (TRACERx, PEACE and CRUK Cancer Immunotherapy Catalyst Network), the CRUK Lung Cancer Centre of Excellence, the Rosetrees Trust, NovoNordisk Foundation (ID16584) and the Breast Cancer Research Foundation (BCRF). This research is supported by a Stand Up To Cancer-LUNGevity-American Lung Association Lung Cancer Interception Dream Team Translational Research Grant (Grant Number: SU2C-AACR-DT23-17). Stand Up To Cancer is a program of the Entertainment Industry Foundation. Research grants are administered by the American Association for Cancer Research, the Scientific Partner of SU2C. CS receives funding from the European Research Council (ERC) under the European Union’s Seventh Framework Programme (FP7/2007-2013) Consolidator Grant (FP7-THESEUS-617844), European Commission ITN (FP7-PloidyNet 607722), an ERC Advanced Grant (PROTEUS) from the European Research Council under the European Union’s Horizon 2020 research and innovation programme (grant agreement No. 835297), and Chromavision from the European Union’s Horizon 2020 research and innovation programme (grant agreement 665233).

### Authors’ contributions

BS carried out all experiments; AP ran the mutation detection algorithms; BS, AP, EN and DS analysed the mutation data; AP and NK analysed public tumour mutation data; DS and CS conceived and coordinated the study; DS wrote the manuscript; all authors helped drafting the manuscript and read and approved the final version.

## Acknowledgements

Not applicable.

## Notes

https://github.com/szutsgroup/platinum_mutagenesis

### Competing Interest Statement

The authors have declared no competing interest.

