## Supplemental figures for "Cisplatin is more mutagenic than carboplatin or oxaliplatin at equitoxic concentrations"

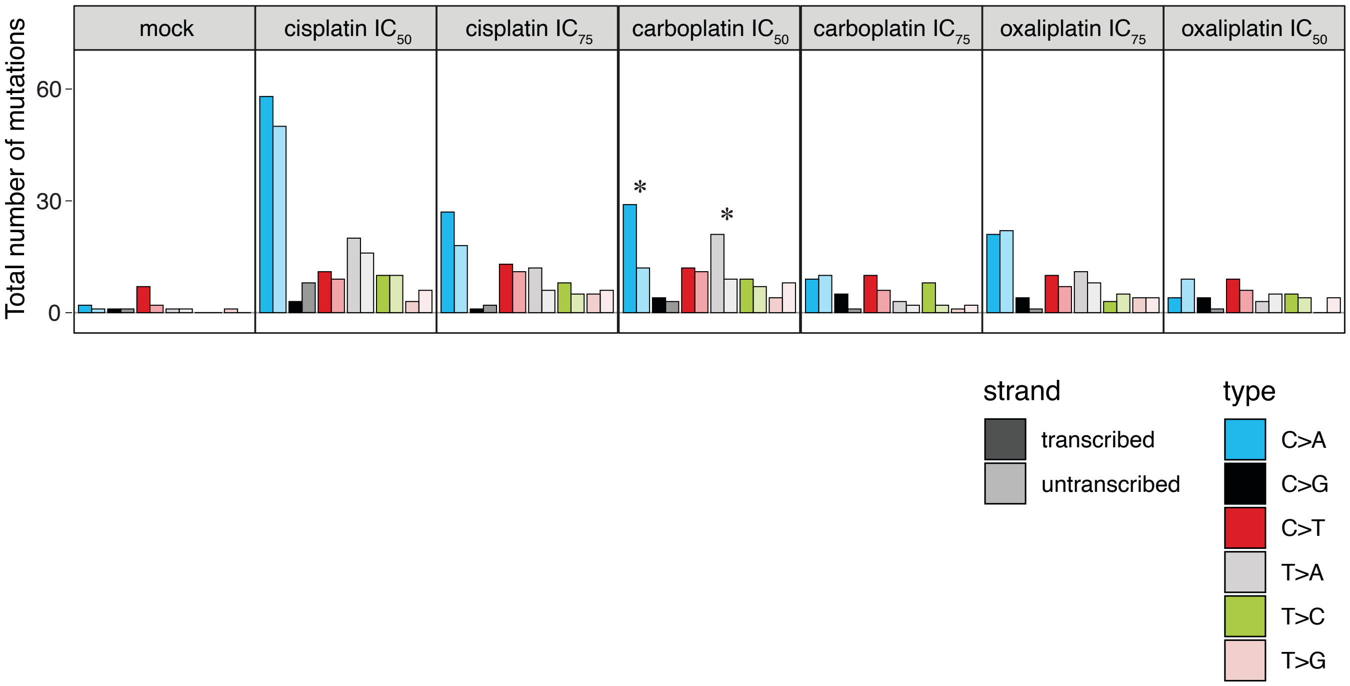


**Supplementary Figure 1.** Transcriptional bias of base substitution mutations.
The total number of genic mutations in five sequenced DT40 cell clones per treatment is shown as indicated (three samples in case of the mock treatment), separated by mutation type and transcriptional strand. Asterisks indicate significant transcriptional strand asymmetries ($P < 0.05$, two-sided Poisson test).

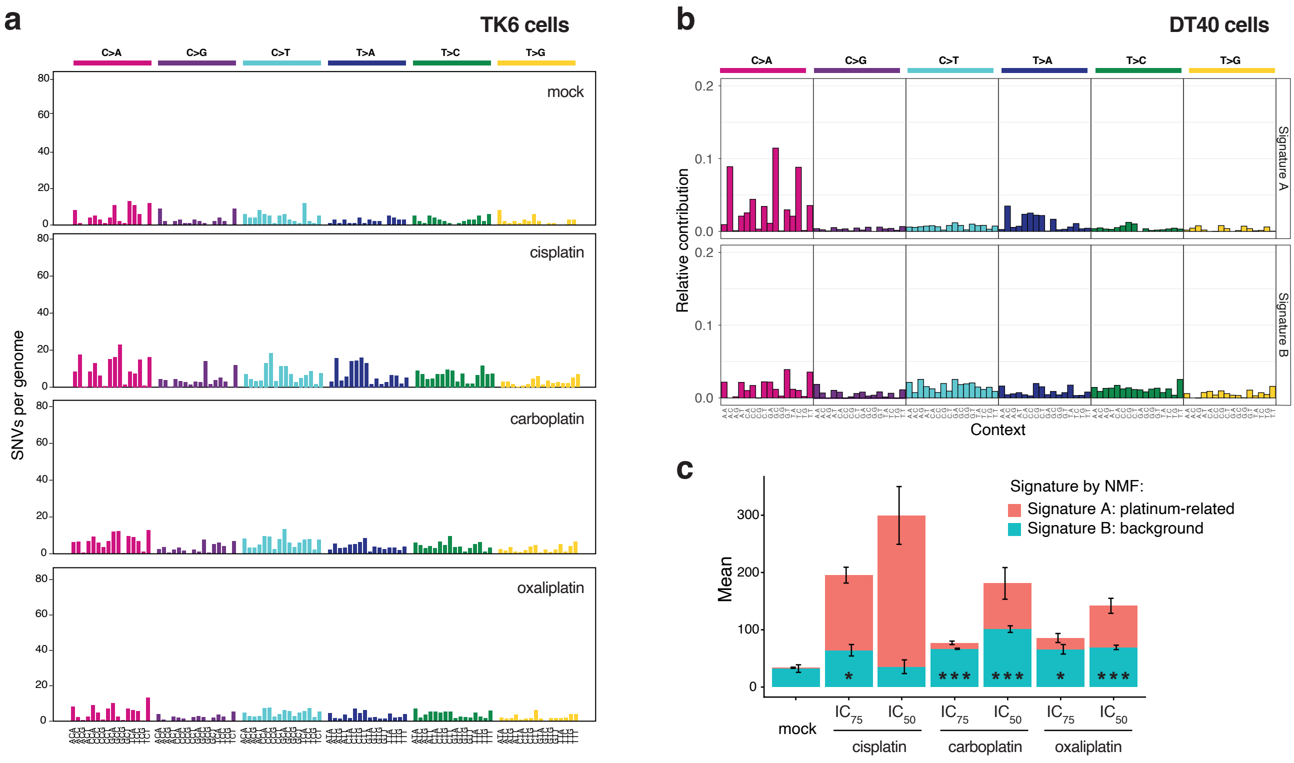


**Supplementary Figure 2.** Analysis of SNV mutation spectra.

(**a**) Base substitution triplet mutation spectra of the mock, cisplatin, carboplatin and oxaliplatin treatments in TK6 cells. The middle base of each triplet, listed at the bottom, mutated as indicated at the top of the panel.

(**b**) De novo NMF of SNV mutation spectra of all sequenced DT40 samples identified two components; triplet mutation spectra of NMF signature A (platinum related) and NMF signature B (background) are shown as the contribution of each triplet mutation type.

(**c**) Mean SNV counts for sequenced DT40 genomes, split using NMF into 'platinum related' and 'background' signatures and then averaged by treatment as indicated. The error bars indicate the S.E.M of the column below, based on five independent treated single cell clones. Significant differences in the number of mutations attributed to the 'background' signature, as compared to the mock treatment, are indicated at the bottom of the columns (unpaired *t*-test, *$P$<0.05, **$P$<0.01, ***$P$<0.001).

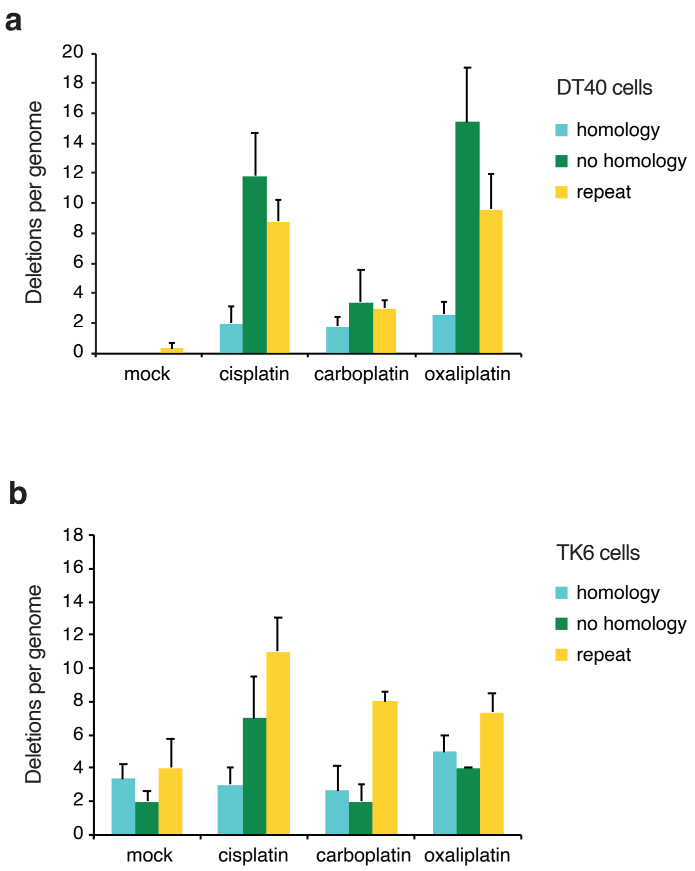


**Supplementary Figure 3.** Short genomic deletions caused by platinum treatment.
A classification of short deletion events by sequence context (homology, no homology and repeat) in DT40 cells (**a**) and in TK6 cells (**b**) treated at $IC_{50}$ concentrations of the respective drugs. The minimum length of classified microhomologies was 1 bp. Error bars indicate S.E.M.

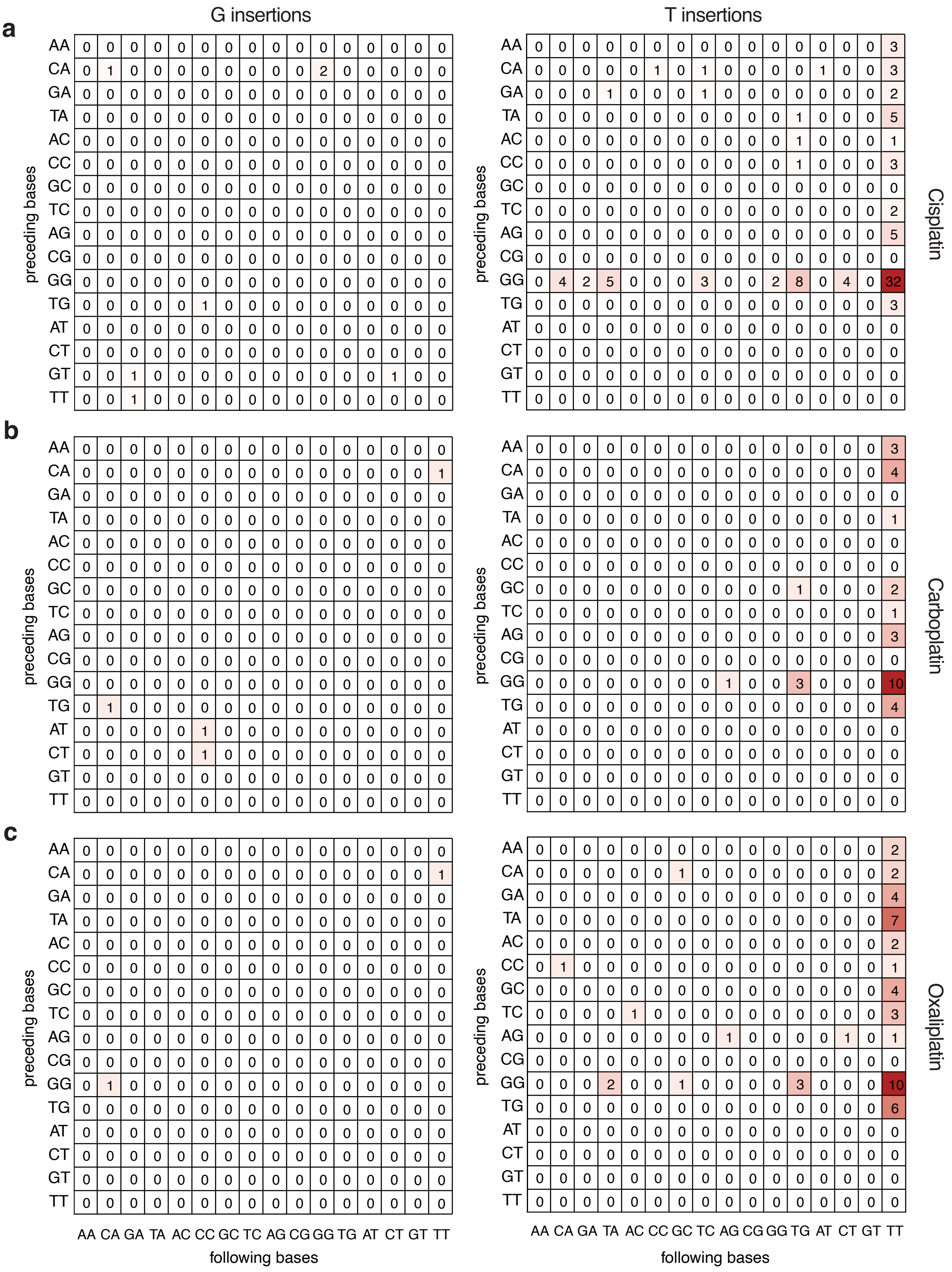


**Supplementary Figure 4.** Sequence context analysis of platinum-induced single nucleotide insertions in DT40 cells.

Heat map of the total number of one nucleotide insertions in all DT40 cell clone genomes treated with each platinum drug (**a** cisplatin, **b** carboplatin, **c** oxaliplatin), classified according to the preceding and the following two bases as indicated. The inserted base is shown above each panel. The equivalent complementary mutations on the two strands are added together and shown as C or T insertions.

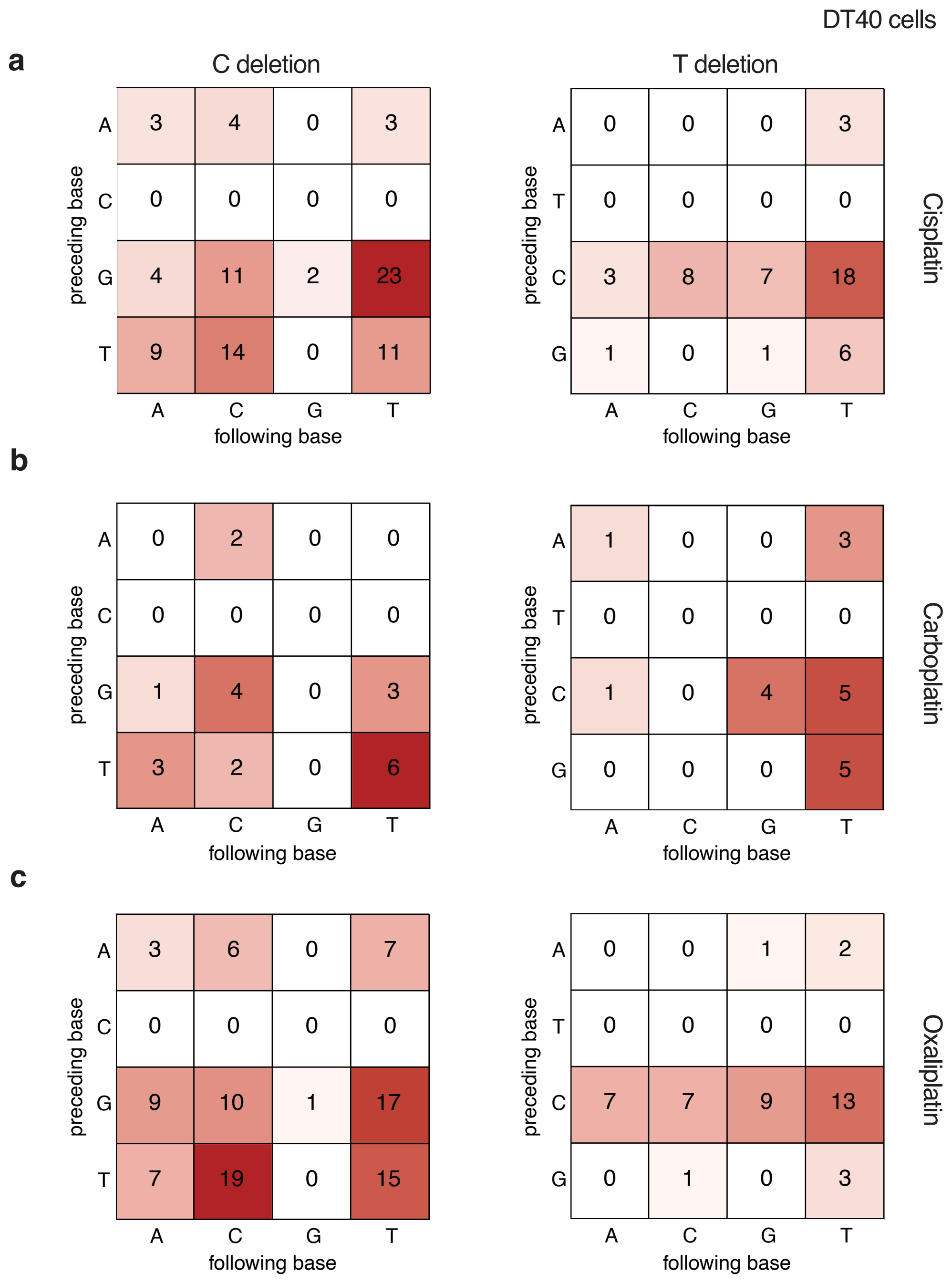


**Supplementary Figure 5.** Sequence context analysis of platinum-induced single nucleotide deletions in DT40 cells.

Heat map of the total number of one nucleotide deletions in all DT40 cell clone genomes treated with each platinum drug (**a** cisplatin, **b** carboplatin, **c** oxaliplatin), classified according to the preceding and the following one base as indicated. The deleted base is shown above each panel. The equivalent complementary mutations on the two strands are added together and shown as C or T deletions.

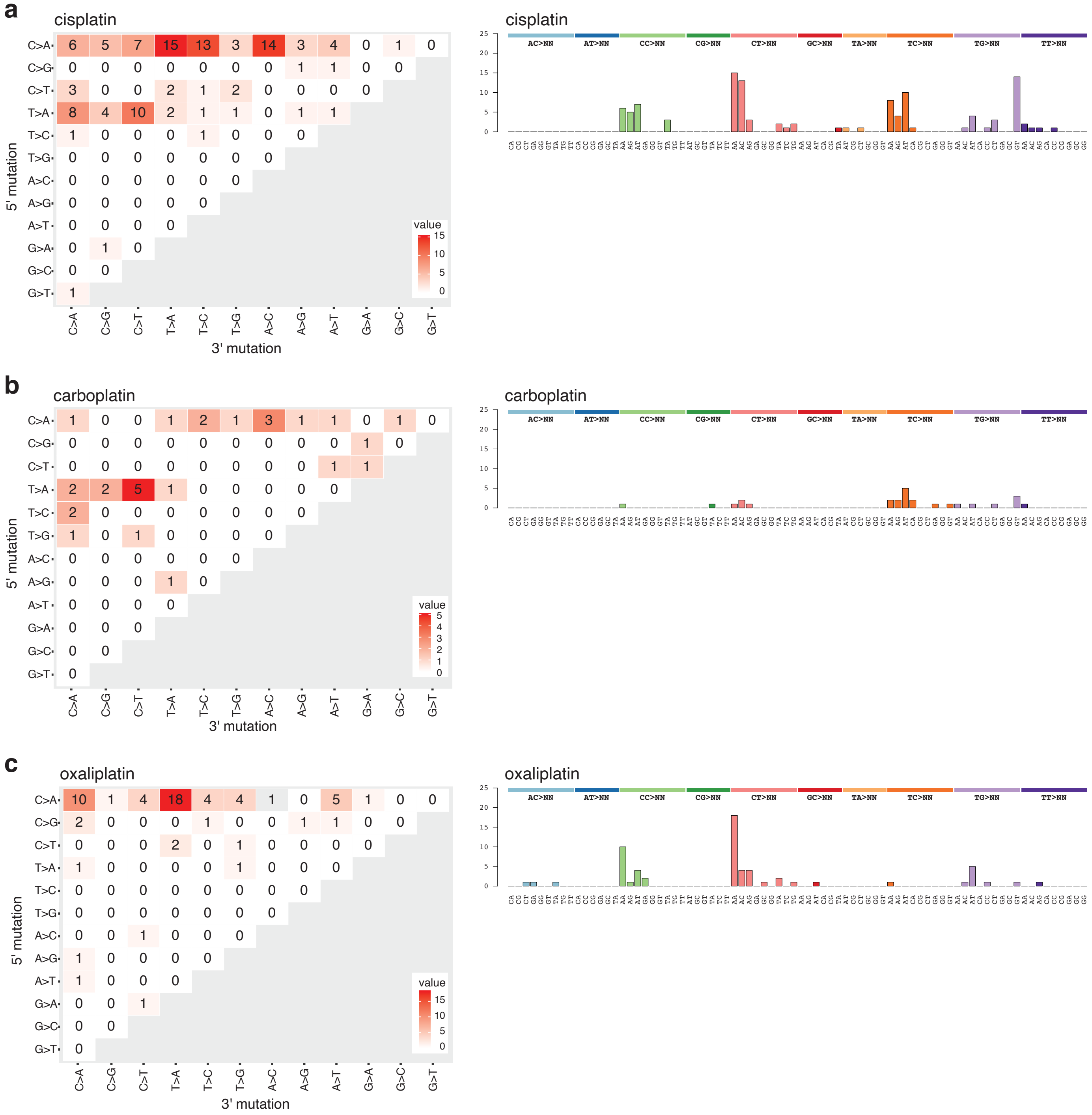


**Supplementary Figure 6.** Sequence analysis of dinucleotide mutations detected in DT40 genomes following cisplatin (**a**), carboplatin (**b**) and oxaliplatin (**c**) treatment. On the left panels, the change in the 5’ base is shown in the rows, while the 3’ base in the columns. The equivalent mutations on the two strands are added together, e.g. GG>TT is shown as CC>AA. The right panels show the same data presented according to the COSMIC double base substitution (DBS) classification.
